# Mitochondrial Genomes of Giant Deers Suggest their Late Survival in Central Europe

**DOI:** 10.1101/014944

**Authors:** Alexander Immel, Dorothée G. Drucker, Tina K. Jahnke, Susanne C. Münzel, Verena J. Schuenemann, Marion Bonazzi, Alexander Herbig, Claus-Joachim Kind, Johannes Krause

## Abstract

The giant deer *Megaloceros giganteus* is among the most fascinating Late Pleistocene Eurasian megafauna that became extinct at the end of the last ice age. Important questions persist regarding its phylogenetic relationship to contemporary taxa and the reasons for its extinction. We analyzed two large ancient cervid bone fragments recovered from cave sites in the Swabian Jura (Baden-Württemberg, Germany) dated to 12,000 years ago. Using hybridization capture in combination with next generation sequencing, we were able to reconstruct nearly complete mitochondrial genomes from both specimens. Both mtDNAs cluster phylogenetically with fallow deer and show high similarity to previously studied partial *Megaloceros giganteus* DNA from Kamyshlov in western Siberia and Killavullen in Ireland. The unexpected presence of *Megaloceros giganteus* in Southern Germany after the Ice Age suggests a later survival in Central Europe than previously proposed. The complete mtDNAs provide strong phylogenetic support for a *Dama-Megaloceros* clade. Furthermore, isotope analyses support an increasing competition between giant deer, red deer, and reindeer after the Last Glacial Maximum, which might have contributed to the extinction of *Megaloceros* in Central Europe.

## Introduction

The extinct giant deer *Megaloceros giganteus* (also Irish Elk), first described by Blumenbach in 1799^1^, stands out amongst the Pleistocene megafauna not only due to its sheer body size, but also because of its immense antlers, which spanned up to 4m in diameter and weighted up to 45kg^2^. Appearing in the fossil record around 400,000 years ago (ya)^3^, its populations are thought to have ranged from Ireland to Lake Baikal^4^. Many theories have been proposed to account for its pattern of distribution across Eurasia in the Late Pleistocene and its extinction in the early Holocene. One unresolved question concerns the reason for the absence of giant deer during the Last Glacial Maximum (LGM, 20,000 – 12,500 ya) in Western and Central Europe, implying that these species had completely withdrawn from the region^5^. Before its purported extinction ca. 6,900 ya in western Siberia, *Megaloceros* recolonized northwestern Europe in the Late Glacial Interstadial^5^; however, no evidence has thus far been found for the presence of giant deer in Southern- and Central Europe.

The phylogenetic position of *Megaloceros* within the family *Cervidae* is still debated. The presence of large palmate antlers in extant fallow deer (*Dama dama*) and *Megaloceros giganteus* suggests a close relationship between the two species^6,7^, whereas postcranial skeletal characters place *Megaloceros* into a group comprising red deer (*Cervus elaphus*), cheetal (*Axis*), and bush antlered deer (*Eucladoceros*)^8^. Previous genetic studies on short regions of mitochondrial DNA (mtDNA) provide evidence for a closer relationship to fallow deer^9,10^; however, the statistical support for the *Dama - Megaloceros* clade is low, likely due to the partial mtDNA regions studied and the limited availability of modern cervid mtDNA for comparison^11,12^. Here we present two reconstructed nearly complete mitochondrial genomes of large cervid bone fragments found in two cave sites in Southern Germany dating to the late glacial period (ca. 12,000 ya). Phylogenetic analyses reveal that both specimens derive from *Megaloceros giganteus*. Comparison against mtDNAs from 44 extant deer species provides furthermore strong support for a *Dama - Megaloceros* mtDNA clade.

## Results and Discussion

### Morphological Analyses and Dating

During an excavation at the Hohlenstein Stadel cave at the Lone Valley (Baden-Württemberg, Southwestern Germany), an accumulation of large cervid bones was discovered including an almost complete atlas, two scapulae and two pelvic fragments, two ribs, a tooth (M3) and six fragments of a tibia. Apart from these finds, a metatarsus shaft fragment from a large cervid was obtained from the Hohle Fels cave at Schelklingen (Baden-Württemberg, Southwestern Germany). We genetically analyzed one of the six tibia fragments (ST/213/203/144, Fig. 1a) recovered from Hohlenstein Stadel cave, dated to 12,175±50 uncal ya (ETH-41223), and the metatarsus fragment (HF/65/100, Fig. 1b) from Hohle Fels cave, which was dated to 12,370±30 uncal ya (MAMS-16557). Radiocarbon ages were calibrated based on IntCal13 curve and calculated using the Calib 7.0 program.

**Figure 1.**
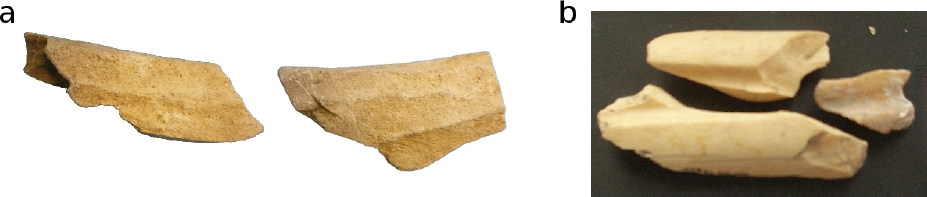
Ancient large cervid remains. (a) A tibia fragment (ST/213/203/144) from the Hohlenstein Stadel cave and (b) a metatarsus fragment (HF/65/100) from the Hohle Fels cave site.

### mtDNA assembly

In order to reconstruct the mitochondrial genomes of our ancient cervid specimens, we isolated the total DNA, turned it into DNA sequencing libraries and enriched for mtDNA using bait generated from modern roe deer with specific primers (table 1). Illumina sequencing on a HiSeq2500 produced 944,648 and 6,123,389 merged reads, for Hohlenstein Stadel and Hohle Fels, respectively. These reads were mapped to the mitochondrial reference sequence of a roe deer (NC_020684.1) and a fallow deer (JN632629.1). Consensus sequences were generated for each sample based on at least 3-fold coverage. Both ancient samples produced identical consensus sequences for shared regions, demonstrating mitochondrial similarity (Supplementary information). However, using fallow deer^13^ as a mapping reference created an almost complete mtDNA sequence (91.52 % for Hohlenstein Stadel and 99.99 % for Hohle Fels), with 7634 unique mapping fragments for the Hohlenstein Stadel sample and 1,009,775 unique mapping fragments for Hohle Fels (table 2). The consensus sequence generated from positions with at least 3-fold coverage from the high-coverage Hohle Fels sample was subsequently used as reference sequence to re-align the Hohlenstein Stadel fragments. Remapping to this consensus sequence provided 7782 mapped fragments after duplicate removal for the Hohlenstein Stadel sample.

**Table 1.**
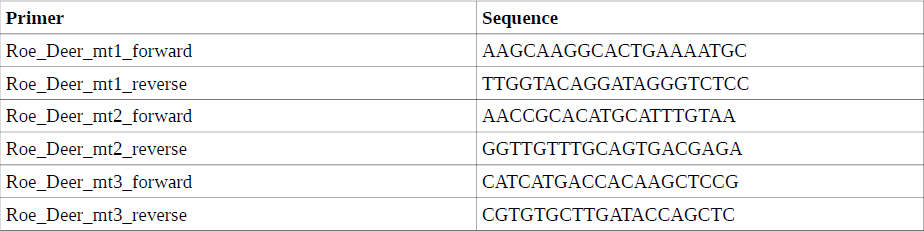
Forward and reverse primer pairs used to generate bait from roe deer mtDNA for targeted mtDNA enrichment.

**Table 2.** Mapping results for the Hohlenstein Stadel sample (ST/213/203/144) and the Hohle Fels sample (HF/65/100). EB: Extraction blank, LB: library blank. Columns from left to right: Sample, calibrated radiocarbon date, number of merged reads, number of unique mapped reads to the fallow deer mtDNA sequence, average mitochondrial genome coverage, average read length, and frequency of C to T substitutions at 5’ end.

| **Sample** | **<sup>14</sup>C cal ya**<br>**2 sigma** | **Total Merged**<br>**Reads** | **Unique mapped**<br>**Reads** | **Unique Average**<br>**Coverage** | **Average Read**<br>**Length** | **%C → T**<br>**substitutions** |
| --- | --- | --- | --- | --- | --- | --- |
| Hohlenstein<br>Stadel | 13904 - 14215 | 944,648 | 7,634 | 28x | 60 | 27 |
| Hohle Fels | 14153 - 14681 | 6,123,389 | 1,009,775 | 5296x | 85 | 49 |
| Hohlenstein<br>Stadel EB | 13904 - 14215 | 422,344 | 193 | 1.3x | 109 | 0 |
| Hohlenstein<br>Stadel LB | 13904 - 14215 | 320,132 | 264 | 1.8x | 109 | 0 |
| Hohle Fels<br>EB | 14153 - 14681 | 50,832 | 48 | 0.2x | 76 | 0 |
| Hohle Fels<br>LB | 14153 - 14681 | 337,962 | 25 | 0.17x | 108 | 0 |

### DNA damage patterns

To authenticate the sequenced fragments as ancient, the frequency of terminal substitutions was analyzed. It has been suggested that C to T substitutions at the 5’ end and G to A substitutions at the 3’end are likely caused by deamination of cytosine causing miscoding lesions; these accumulate over time, and hence are characteristic of ancient DNA^14^. DNA fragments with frequencies of at least 20% for both types of substitutions at the 5’ and 3’ end can be regarded as authentic ancient DNA^15^. We observed a substitution frequency of 27% for C to T changes at the 5' ends and a substitution frequency of 25 % of G to A at the 3' ends of sequence reads in the Hohlenstein Stadel sample (Fig. 2a). In the Hohle Fels sample 49% of the sequence reads showed a C to T substitution at the 5’ ends and 48% a G to A substitution at the 3' ends (Fig. 2b). These values almost reach the theoretical maximum of 50% deamination at single stranded overhangs with a double strand library preparation protocol and indicate authentic ancient mtDNA in both faunal remains.

**Figure 2.**
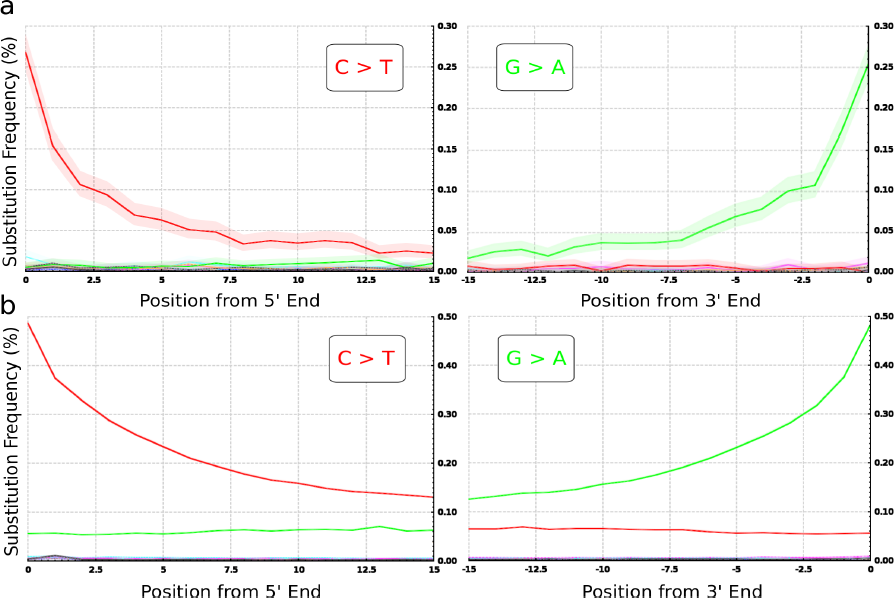
Substitution pattern at the 5' and 3' ends of the aligned sequence reads from the Hohlenstein Stadel sample (a) and the Hohle Fels sample (b). The misincorporation plots were generated using a custom software extension package (Krause J. *et al*. A complete mtDNA Genome of an Early Modern Human from Kostenki, Russia. *Curr. Biol*. **20**, 231-236 (2010)).

### Phylogenetic Analysis

The reconstructed mtDNAs from both ancient cervid bones were aligned with 44 publicly available full mitochondrial genomes of extant cervids. The hypervariable D-loop region was excluded from the cervid mtDNA alignment due to its fast evolutionary rate that may decrease the phylogenetic resolution. We used MEGA 6.0.6 to construct both a maximum-likelihood (ML) tree (Fig. 3a) and a maximum-parsimony (MP) tree (Fig. 3b) and robustness of both methods was tested with 1000 bootstrap replicates. Both tree reconstructions were based on a total of 14,147 positions. Alpine musk deer (*Moschus chrysogaster*, KC425457.1) was chosen as an outgroup. Both ML and MP topologies place the mtDNA sequences reconstructed from our two ancient cervid bones (Hohlenstein Stadel and Hohle Fels) in a completely resolved clade with both extant fallow deer subspecies (*Dama dama* and *Dama mesopotamica*), and exclude them from the group comprising Pere David's Deer (*Elaphurus davidianus*), *Rusa* sp. and red deer (*Cervus elaphus* sp). The other large cervid present in Europe during the Pleistocene, the European elk, *Alces alces*, can also be excluded as the source of our ancient cervid bones.

**Figure 3.**
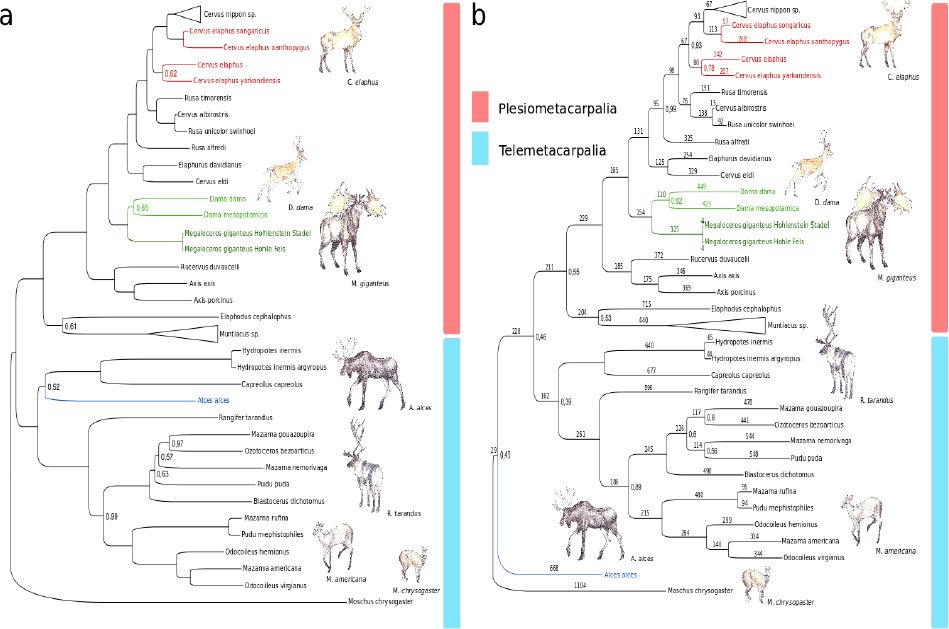
Phylogenetic trees of full mtDNA sequences from 44 extant cervid species and two ancient mtDNA sequences from two ancient cervid bones likely representing *Megaloceros giganteus*. Each tree is based on 14,147 positions. Bootstraping was performed with 1000 bootstrap replicates. Only bootstrap values different from 1 (=100) are indicated at inner nodes. (a) Maximum-likelihood tree. (b) Maximum-Parsimony tree. Both topologies place *Megaloceros giganteus* together with fallow deer (*Dama* sp.) in a basal position to the red deer clade (*Cervus elaphus*). Both trees were rooted with musk deer (*Moschus chrysogaster*) as outgroup. Deer drawings were kindly prepared and provided by Kerttu Majander.

The relationship of our ancient cervids to both *Dama* species within the *Dama* clade is, however, not completely resolved. As *Dama* was only introduced to Europe in the Medieval period^16^ and is absent in the palaeontological record of Western Eurasia, and since both ancient cervid bones derive morphologically from a large cervid, which is not elk (*Alces alces*) based on phylogenetic evidence, we conclude that both specimens originate from *Megaloceros giganteus*. To test this hypothesis we further compared our reconstructed ancient cervid mtDNAs to cytochrome b (*cytb*) regions of *Megaloceros* mtDNA previously published (table 3). We find 17 and 1 nucleotide differences between the Hohlenstein Stadel sample and the Hohle Fels sample, respectively, compared to the previously published *cytb* sequence from a complete *M. giganteus* skeleton from the Kamyshlov site in western Siberia^9^ and 19 and 1 nucleotide differences compared to the *cytb* sequence obtained from a *M. giganteus* astralagus from Killavullen in Ireland^10^, whereas for the same region we find 101 (100) differences to fallow deer^9^ and 103 (95) differences to red deer^17^. The small nucleotide distance of our ancient large cervids and both previously determined giant deer *cytb* sequences confirm the attribution of our samples to *Megaloceros giganteus*. The close genetic relationship between the large cervid bone from Hohle Fels and the giant deer skeleton from Kamyshlov in western Siberia (AM072744.1)^9^ and the giant deer astragalus from Killavullen in Ireland (AM182645.1)^10^, respectively, suggests furthermore a close maternal relationship and low genetic diversity between Nothwestern European, Eastern and Central European giant deer populations.

**Table 3.** Nucleotide sequence distances between the reconstructed and previously published cervid *cytb* sequences.

| Sequence | *M. giganteus cytb*<br>(AM072744.1)<sup>9</sup> | *M. giganteus cytb*<br>(AM182645.1)<sup>10</sup> | Fallow Deer *cytb*<br>(AJ000022.1)<sup>9</sup> | Red Deer *cytb*<br>(AB924664.1)<sup>17</sup> |
| --- | --- | --- | --- | --- |
| Hohlenstein Stadel *cytb* | 17 | 19 | 101 | 103 |
| Hohle Fels *cytb* | 1 | 1 | 100 | 95 |

### Stable Isotopes

To evaluate the stable isotope signature of our ancient cervids from the Hohle Fels and Hohlenstein Stadel cave sites, stable isotopes from collagen carbon (^13^C) and nitrogen (^15^N) were measured and compared to large cervids present in Central Europe during the Pleistocene such as red deer, reindeer, and giant deer^18,19,20^. Pre-LGM (ca. 35,000 uncal ya) *Megaloceros* samples from Southern France and Belgium typically show isotopic signatures of collagen comparable to those of red deer (*Cervus elaphus*), while reindeer (*Rangifer tarandus*) provides systematically higher δ^13^C_coll_ values likely due to the consumption of lichen^18^ (Fig. 4a). During the Late Glacial period (13,000 – 12,000 uncal ya), the ^13^C-based distinction among larger cervids from the Swabian, French, and Swiss Jura decreases. Stable isotope signatures of our both ancient cervid bones from Hohle Fels cave and Holenstein Stadel cave fall inside the red deer-reindeer cluster reflecting a potential overlap in diet and habitat (Fig. 4b).

**Figure 4.**
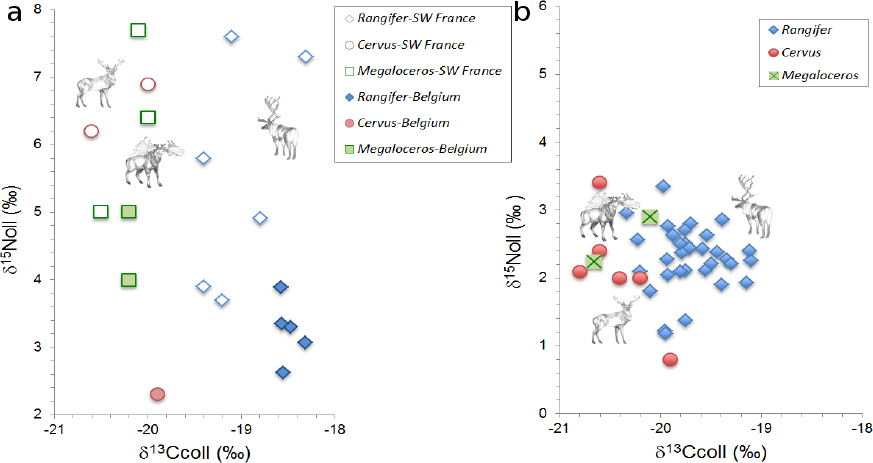
(a) Stable isotope values of reindeer (*Rangifer*), red deer (*Cervus*) and *Megaloceros* before the LGM in SW France and Belgium and (b) after the LGM in the Swabian, Swiss and French Jura indicate a decrease in the distinction between the isotope signatures of the three cervid species after the LGM, which might be due to overlapping diet and habitat. The deer drawings were kindly prepared and provided by Kerttu Majander.

### Discussion

We obtained nearly complete mtDNA sequences from two ancient cervid bones from the Swabian Alb dated to 12,175±50 uncal ya (13,904 – 14,215 cal ya) and 12,370±30 uncal ya (14,153 – 14,681 cal ya), respectively. Both sequences are distinct from 44 mtDNAs of extant cervids. The phylogenetic analyses suggests that the reconstructed mtDNAs are maternally closely related to fallow deer (*Dama*). Based on the phylogenetic position of our reconstructed ancient mtDNAs, their close genetic relationship to the previously determined partial *cytb* sequence from a complete giant deer skeleton from western Siberia^9^ and to the complete *cytb* sequence from a giant deer astralagus from Ireland^10^ and due to the absence of fallow deer in Europe in the Pleistocene as well as due to the size of the bones, we conclude that both specimens derive from *Megaloceros giganteus*. We find strong support for a close maternal relationship to both *Dama* species. The maternal relationship within the *Megaloceros-Dama* clade however could not be resolved in our phylogeny suggesting an almost equal genetic distance of the two fallow deer species and giant deer. Our results disagree with the morphological conclusions that *Megaloceros* is closer related with a group comprising *Cervus*, *Axis*, and *Eucladoceros*^8,21^, and that the occurrence of palmate antlers in *Megaloceros* and *Dama* must be the result of homoplasy. Our results also disagree with the conclusions derived from short mtDNA sequences such as partial *cytb* reported by Kuehn and colleagues^22^ which suggested a *Cervus-Megaloceros* clade and attributed the palmate antlers in *Megaloceros* and *Dama* to homoplasy. Instead, our results support the hypothesis of a *Megaloceros-Dama* clade suggested by previous studies based on morphological features and phylogenetic analyses from *cytb*^2,3,9,10^.

*Megaloceros* is traditionally considered a species adapted to open-areas that could have suffered from the development of forest in the early Holocene, most likely because the huge antlers would have restricted the movement of males in dense woodland^5^. However, anatomy and distribution suggest that *Megaloceros* was a mixed feeder^5^, and carbon-13 results on enamel of *Megaloceros* of previous interglacial periods support the possibility of a boreal habitat^23^. Isotope signatures of our two *Megaloceros* samples reveal that the diversity in habitat and diet decreased after the LGM between *Megaloceros*, red deer, and reindeer, probably resulting in an increased competition among the deer species in Central Europe. The overlapping niches and thus the increased competition with other deer species could explain, at least in part, the local extinction of *Megaloceros* in Southern Germany.

In conclusion, we generated two almost complete mitochondrial genomes from large cervid bones from the Hohle Fels and Hohlenstein Stadel caves in Southwestern Germany that date back to 12,000 uncal ya. Phylogenetic comparisons to contemporary deer mtDNA and previously determined ancient *cytb* DNA suggest that both mtDNA genomes derive from *Megaloceros giganteus*, which demonstrates its presence in Central Europe after the LGM. The close maternal relationship with the two fallow deer species resulting in a near polytomy in the *Dama-Megaloceros* clade questions the morphology-based grouping of giant deer and fallow deer in two separate genera. To date there has been no evidence that *Megaloceros* recolonized Central Europe after the LGM^5^; our findings provide support that *Megaloceros* returned to central parts of Europe even if the presence of humans might have hindered the re-colonization. In addition stable isotopes from our ancient cervid bones suggest a direct competition with other cervid species at the onset of the Holocene, potentially due to the lack of niche partitioning. Thus enviromental factors may have played an important role in the final extinction of the giant deer.

## Material and Methods

### Extraction of ancient DNA

Bone samples were exposed to UV-light overnight to remove surface contamination. A sample of 50mg was removed from the inside of the longbone of each bone using a dentistry drill. DNA extraction was carried out using a guanidinium-silica based method^24^. For each sample a DNA library was prepared according to published protocols^25^. Sample-specific indexes were added to both library adapters to allow differentiation between individual samples after pooling and multiplex sequencing^26^. Indexed libraries were amplified in 100µl reactions followed by purification over Qiagen MinElute spin columns (Quiagen, Hilden, Germany) and quantification using Agilent 2100 Bioanalyzer DNA 1000 chip. Target enrichment of mitochondrial DNA was performed by capture of the pooled libraries using bait generated from modern roe deer (*Capreolus capreolus)* mitochondrial DNA^27^. The bait was generated by use of three primer sets (table 1) designed with the Primer3Plus software package. All extractions and pre-amplification steps of the library preparation were performed in clean room facilities and one negative control was included for each reaction.

### Sequence Processing, Assembly, Duplicate Removal

Library pools were sequenced on the Illumina Hiseq 2500 platform using two index reads (2*100+7+7 cycles) following the manufacturer's protocol. De-indexing was performed by sorting all sequences corresponding to their p7 and p5 combinations using the CASAVA software version 1.8. Forward and reverse reads were merged into single sequences if they overlapped by at least 11bp^28^. Unmerged reads were discarded and merged reads were filtered for a length of at least 30bp. Mapping of the length-filtered reads and removal of duplicate reads was performed using a custom mapping iterative assembler (MIA) which was developed to take into account sequence errors which commonly occur from ancient DNA damage^29^. Reads were mapped to a full mitochondrial genome reference sequence of the roe deer, *Capreolus capreolus* (NC_020684.1). To achieve a higher resolution in the topology, in a second round sequence reads were mapped to a full mitochondrial genome of the fallow deer *Dama dama* (JN632629.1), and in a third round to the consensus sequence of our ancient putative *Megaloceros giganteus* sample HF/65/100, which was generated by mapping to the *Dama dama* mitochondrial reference sequence as described.

### Analysis of ancient DNA damage patterns

C to T and G to A substitution patterns were obtained from the sequences using a custom software developed as an extension package to handle the output format of the mapping iterative assembler^29^.

### Multiple Sequence Alignment and Molecular Phylogenetic Analyses

A multiple sequence alignment was generated from 44 full mitochondrial genome Genbank^30^ sequences of extant cervid taxa together with the assembled mitochondrial genome sequences of our putative *Megaloceros* samples using ClustalW (Larkin, M.A. *et al*. ClustalW and ClustalX version 2. *Bioinformatics* **23**: 2947-2948 (2007). Alignments and phylogeny constructions were conducted in MEGA 6.0.6. The mitochondrial D-Loop region was excluded using the BioEdit sequence alignment editor (Hall, T.A. BioEdit: a user-friendly biological sequence alignment editor and analysis program for Windows 95/98/NT. *Nucleic Acids Symp. Ser*. **41**:95-98 (1999)) and phylogenies were constructed from a total of 14,147 positions. Both maximum-likelihood and maximum-parsimony topologies were generated for all positions for which coverage was at least three-fold in each of the ancient sequences. Alignment columns with gaps or missing data were eliminated. Bootstrap support values were obtained over 1000 replicate data sets, using alpine musk deer as an outgroup (*Moschus chrysogaster*, KC425457.1). The phylogenetic trees were edited in FigTree version 1.4.0 (*http://tree.bio.ed.ac.uk/software/figtree*).

### Pairwise Comparison of Cytochrome b Sequence differences

To identify cytochrome b (*cytb*) coordinates within our reconstructed mitochondrial sequences, these were aligned to previously published *cytb* sequences of *Megaloceros giganteus* (AM072744.1, AM182645.1), fallow deer (AJ000022.1), and red deer (AB924664.1) using MEGA 6.0.6 and sequences outside of the aligned regions were discarded. Nucleotide sequence distances were then calculated by pairwise alignment between each of our ancient *cytb* sequences and each of the previously published *cytb* sequences using BLAST.

### Stable Isotope Analyses

To study the habitat pattern revealed by stable isotopes, collagen was extracted from both bone fragments and carbon (^13^C) and nitrogen (^15^N) were measured. The results were combined with stable isotope data obtained from ancient reindeer (*Rangifer tarandus*) and red deer (*Cervus elaphus*) remains of the Swabian, French, and Swiss Jura dating 13,000 to 12,000 uncal year ago, which corresponds roughly to the GI-1e interstadial^18,19,20^. The results were compared to stable isotopes from morphologically defined deer specimens including giant deer (*Megaloceros giganteus*) dating to the pre-LGM from southwestern France (SW France) and Belgium^18,31^.

Isotopic analysis (δ^13^C_coll_, δ^15^N_coll_) was conducted at the Department of Geosciences of Tübingen University using a Thermo Quest Delta+XL mass spectrometer coupled to a NC2500 CHN-elemental analyzer, which provides elemental analysis (Ccoll, Ncoll). The international standards used include marine carbonate (V-PDB) for δ^13^C and atmospheric nitrogen (AIR) for δ^15^N. Analytical error, based on within-run replicate measurement of laboratory standards (albumen, modern collagen, USGS 24, IAEA 305A), was ±0.1‰ for δ^13^C values and ±0.2‰ for δ^15^N values. Reliability of carbon and nitrogen isotopic values was established by measuring the chemical composition, with C/N_coll_ atomic ratio within the range of 2.9 to 3.6^32^.

## ACKNOWLEDGEMENTS

We thank Kerttu Majander for the elaborate deer drawings and Kirsten Bos for valuable comments that greatly improved the manuscript. This research was funded by European Research Council Starting Grant APGREID (to A.I. and J.K.).

### Author Contributions

A.I. and J.K. conceived and designed the research. T.K.J., S.C.M. and C.J.K. provided the bone samples and conducted initial morphological analyes. M.B. and V.J.S performed the extraction of mtDNA and preparation of sequencing libraries. A.I. and A.H. performed the bioinformatic analyses and D.G.D. conducted the stable isotope analyses. A.I. wrote the manuscript and J.K. and D.G.D mostly contributed to the discussion. All authors reviewed the manuscript.

### Competing financial interests

The authors declare no competing financial interests.

